# CD11c^+^CD88^+^CD317^+^ myeloid cells are critical mediators of persistent CNS autoimmunity

**DOI:** 10.1101/2020.06.10.144956

**Authors:** Navid Manouchehri, Rehana Z. Hussain, Petra D. Cravens, Brian T. Edelson, Gregory F. Wu, Anne H. Cross, Richard Doelger, Nicolas Loof, Todd N. Eagar, Thomas G. Forsthuber, Laurent Calvier, Joachim Herz, Olaf Stüve

## Abstract

**Background:** Natalizumab, a humanized monoclonal antibody (mAb) against α4-integrin, reduces the number of dendritic cells (DC) in cerebral perivascular spaces in multiple sclerosis (MS). Selective deletion of α4-integrin in CD11c^+^ cells should curtail their migration to the CNS and ameliorate experimental autoimmune encephalomyelitis (EAE).

**Methods:** We generated CD11c.Cre^+/-^*ITGA4*^fl/fl^ C57/Bl6 mice to selectively delete α4-integrin in CD11c^+^ cells. Active immunization and adoptive transfer EAE models were employed. Multi-parameter flow cytometry was utilized to immunophenotype leukocytes. Single-cell RNA sequencing (scRNA-seq) was used to profile individual cells.

**Results:** α4-integrin expression by CD11c^+^ cells was significantly reduced in primary and secondary lymphoid organs in CD11c.Cre^+/-^*ITGA4*^fl/fl^ mice. In active EAE, a delayed disease onset was observed in CD11c.Cre^+/-^*ITGA4*^fl/fl^ mice, during which CD11c^+^CD88^+^ cells were sequestered in the blood. Upon EAE onset, CD11c^+^CD88^+^ cells accumulated in the CNS and expressed CD317^+^. In adoptive transfer experiments, CD11c.Cre^+/-^*ITGA4*^fl/fl^ mice had ameliorated clinical disease associated with diminished numbers of CNS CD11c^+^CD88^+^CD317^+^ cells. The transcription profile of CD11c^+^CD88^+^CD317^+^ cells placed them within previously defined microglia-like cells in human CSF. We show that activated, but not naïve microglia expressed CD11c, CD88, and CD317. Finally, anti-CD317 treatment prior to clinical EAE substantially enhanced recovery.

**Conclusion:** CD11c^+^CD88^+^CD317^+^ cells in the CNS promote inflammatory damage. Transcriptional analysis identifies CD11c^+^CD88^+^CD317^+^ cells as a unique myeloid subset in human CSF. The disease-propagating effects of these cells can be antagonized using anti-CD317 mAb.

## Introduction

Autoimmune disorders of the central nervous system (CNS), including multiple sclerosis (MS), are thought to be mediated by aberrant adaptive immune responses against self-antigens. In experimental autoimmune encephalomyelitis (EAE), a model of MS, activated myelin-reactive CD4^+^ T helper cells are the main drivers of disease activity ^1^. In active EAE, dendritic cells (DC) are professional antigen presenting cells (APC) at the inoculation site, namely in draining lymph nodes and other secondary lymphoid organs, where they present myelin autoantigen to naïve CD4^+^ T lymphocytes^2^. Myeloid APC within the CNS are essential for the re-activation and retention of these autoreactive CD4^+^ T cells, and for perpetuation of disease activity. Specifically, myeloid cells within cerebral perivascular spaces previously considered DC based solely on their expression of CD11c are sufficient to permit EAE ^3^. Myeloid APC, including DC, use α4-integrin to gain access to sites of ongoing inflammation. We previously showed that the number of DC was significantly reduced in cerebral perivascular spaces in autopsy material of an MS patient treated with natalizumab, a humanized monoclonal antibody against α4-integrin ^4^. Conceivably, antagonizing or diminishing the function of α4-integrin selectively in myeloid cells should ameliorate EAE through an impaired reactivation of CNS-specific CD4^+^ T cells and a reduction of direct inflammatory effects exerted by these cells.

To further understand the role of CD11c^+^ cells within the CNS during CD4^+^ T cell-mediated CNS autoimmunity, we generated and characterized CD11c.Cre^+/-^*ITGA4*^fl/fl^ mice, which lack α4-integrin expression in CD11c^+^ cells. We identified CD11c^+^CD88^+^CD317^+^ as mediators of persistent clinical EAE, and propagators of inflammation within the CNS.

## Materials and Methods

### Generation of CD11c.Cre^+/-^ α4 integrin deficient mice

Female homozygote *ITGA4-floxed* (*ITGA4*^fl/fl^) C57/BL6 mice, generated by Dr. Thalia Papayannopoulou, University of Washington ^5^ were crossed with commercially available CD11c/Cre^+^ C57/BL6 males (jax.org; Stock No: 008068) ^6^, and the progeny were backcrossed to generate CD11c^+/-^*ITGA4*^fl/fl^ mice. These mice were extensively characterized ^7^. Each generation of mice was genotyped via genomic polymerase chain reaction (PCR) using CD11c-specific primers. C57BL/6 wild type (WT) mice were purchased from the Jackson Laboratories, Bar Harbor, ME. C57BL/6 WT mice were bred and maintained in a pathogen-free facility at the University of Texas Southwestern Medical Center (UTSW) according to guidelines provided by the National Institute of Health and our institution. Male and female mice were utilized for experiments. We observed no differences regarding disease scores, cellular composition, or any of the biochemical and cellular outcomes between the two sexes (data not shown). The UTSW Institutional Animal Care and Use Committee (IACUC) approved all experiments and procedures.

### Peptides

Mouse myelin oligodendrocyte glycoprotein peptide 35-55 (MOG_p35-55_) (MEVGWYRSPFSRVVHLYRNGK), synthesized by Fmoc chemistry by Quality Controlled Biochemicals, Inc. (QCB; Hopkinton, MA) and CS Bio (Menlo Park, CA), was utilized for active immunization EAE.

### Experimental autoimmune encephalomyelitis

To induce active EAE, experimental mice aged 8-12 weeks were immunized subcutaneously with myelin MOG_p35-55_ (100 μg/100 μl/mouse), emulsified in an equal volume of Complete Freund Adjuvant (CFA) containing 4 mg/mL H37Ra *M. Tuberculosis* (Difco, BD, Franklin Lakes, NJ) in each flank as described ^8^. Upon immunization and 48-hours later, experimental animals received intra-peritoneal injection of 200 ng PTX in 200 μL PBS.

For adoptive transfer EAE, donor cells were harvested from the draining axillary and inguinal lymph nodes (LN) of female WT C57BL/6 mice 10 days after active immunization with MOG_p35-55_ as described above and single cell suspensions were prepared. CD4^+^ T cells, including MOGp35-55-reactive T cells, were purified via negative selection (EasySep™ Mouse CD4^+^ T cell Isolation Kit, STEMCELL Technologies, Lot: 18H94653). Purified CD4^+^ T cells were cultured for 72 hours with interleukin-12 (IL-12) and MOG_p35-55_. Subsequently, cultured cells were injected to recipient mice (10=10^6^ cells/mouse i.p).

For treatment experiments, mice were injected with 250 μg of a rat anti-mouse IgG2bκ anti-CD317 mAb (BioxCell, clone 927) i.p. on days 7 and 8 post immunization.

All experimental animals were observed at least twice daily and disease severity scores were recorded based on a standard EAE scoring system; briefly: 0 = no observable clinical signs, 1 = loss of tail tone or mild hid limb weakness but not both, 2 = limb tail and mild hind limb weakness, 3 = moderate hind limb weakness with or without unilateral hind limb paralysis, 4 = complete bilateral hind limb paralysis, 5 = bilateral hind limb paralysis with fore limb weakness or moribund state or death. The following cumulative interventions were performed for all experiment animals. Moist chow was provided daily once the animal reached clinical score of 3. Regular palpation of the bladder and manual assistance with urination was performed for animals with disease score of 4; animals with EAE score of 4 were kept separated from cage mates with lower scores. Persistence of disease severity at EAE score 4, for at least for 72 hours or observation of EAE score 5 regardless of initiation time warranted euthanasia using carbon dioxide asphyxiation followed by cervical dislocation as the secondary physical method.

### Isolation of lymph node cells and splenocytes

Cells from lymph nodes and spleen were isolated by pressing through a 70 μm nylon mesh cell retainer as described ^8–10^. Cells were treated with red blood cell (RBC) lysis buffer (Sigma-Aldrich, St. Louis, MO, USA), washed twice with cold phosphate buffered saline (PBS), and re-suspended in PBS for counting by hemocytometer.

### Bone marrow cell isolation

Bone marrow cells were isolated from the femurs and tibias of mice as described ^10^. Bones were crushed with a mortar and pestle. Specimens were then passed through a 70 μm nylon mesh cell retainer. Cells were treated with RBC lysis buffer (Sigma-Aldrich, St. Louis, MO, USA), washed 2 times with cold PBS, and re-suspended in PBS for counting with hemocytometer.

### Enzymatic CNS digestion

Mice were anesthetized with 250 mg/kg i.p. injection of tribromoethanol (Avertin, Sigma Aldrich, St Louis, MO) solution and perfused with PBS through the left ventricle as described ^8^. Brains were dissected from the skull and spinal cords were flushed from the spinal column with PBS. Tissues were finely minced using a sterile scalpel and homogenized in cold PBS. We used a commercially available neural tissue dissociation kit for enzymatic dissociation following the manufacturer’s protocol (Miltenyi Biotec, San Diego, CA, USA). Specimens were subsequently washed with cold PBS. 37% Percoll^™^ PLUS (GE Healthcare) gradient was used to remove the residual myelin. The myelin-free single cell suspensions were counted with hemocytometer.

### Separation of blood leukocytes

Submandibular bleeding of actively immunized mice was performed on day 12 post immunization. Approximately, 400 μl of peripheral blood was collected in EDTA tubes from each mouse. The samples from CD11c.Cre^+/-^*ITGA4*^fl/fl^ mice and WT controls were pooled separately. The pooled samples were diluted with 1:1 ratio with PBS and slowly added on top of 100% Ficoll® in a 5 ml cryo tube. Ficoll® gradient layering allowed to separate peripheral blood mononuclear cells (PBMC) without the need for RBC lysis. The collected cells were counted by hemocytometer.

### *In vitro* PBMC culture studies

A total of 0.5 x 10^6^ PBMC were plated per well in 48 well plates in culture medium consisted of Roswell Park Memorial Institute (RPMI) 1640 (Invitrogen) supplemented with L-glutamine (2 mmol/L), sodium pyruvate (1 mmol/L), non-essential amino acids (0.1 mmol/L), penicillin (100 U/mL), streptomycin (0.1 mg/mL), 2-mercaptoethanol (1 μL of stock), and 10% fetal bovine serum (FBS). 10 ng/mL IL-4, and 50 ng/ml granulocyte-monocyte colony stimulating factor (GM-CSF), with or without 20 ng/mL of interferon gamma (IFNγ) was added to wells. The culture medium was changed every 48 hours, and cells were harvested at 72 hours for flow cytometry analysis.

### Multi parameter flow cytometry

A total of 1 x 10^6^ mononuclear cells suspended in PBS were stained using fixable viability dye (Zombie NIR-Biolegend), and incubated for 15 minutes at room temperature following manufacturer’s recommendation. Cells were then washed using 2% FBS-PBS (FACs buffer) at 1500 rpm for 5 minutes, followed by incubation with 1 μg Fc Block (anti-CD16/32, Tonbo Biosciences) for 15 minutes at 4°C. Cells were then stained with the following antibodies: CD45 (Alexa Flour 700-Biolegend, Clone: 30-F11), CD11b (V450-BD Horizon, Clone: M1/70), CD19 (BV510-Biolegend, Clone: 6D5), CD3 (V500-BD Horizon, Clone: 500A2), CD11c (APC-Biolegend, Clone: N418), MHCII (BV785, Biolegend, Clone: M5/114.15.2), CD49d (α4-integrin; FITC, Biolegend, Clone: R1-2), CD88 (PE, Bilegend, Clone: 20/70), CD317 (BV421, Biolegend, Clone: 927) CD26 (FITC, Biolegend, Clone: H194-112) for 45 minutes at 4°C. Events were recorded via BD FACS LSR Fortessa (The Moody Foundation Flow Cytometry Facility, UT Southwestern), equipped with Diva acquisition software (BD Bioscience). Cells were gated according to morphology side scatter (SSC-A) vs forward scatter (FSC-A). Doublets were excluded (FSC-A vs FSC-H and SSC-A vs SSC-H). Live cells were selected using the viability dye. The complete gating strategy is presented in **Supplementary Figure 1**. In each sample a minimum of 50 x 10^3^ live events were recorded. FlowJo software (BD Bioscience) was used for data analysis.

### T cell proliferation assay

To determine the capability of APCs to present CD4^+^ T cells with auto antigens, flow cytometry proliferation assays using the fluorescent dye carboxyfluorescein succinimidyl ester (CFSE) were performed. Ten days post active immunization, spleens and lymph nodes from CD11c.Cre^+/-^*ITGA4*^fl/fl^ mice and WT controls were harvested and processed into single cell suspension as described above. Splenocytes and lymphocytes were stained with CFSE and plated at 1 x 10^6^ cells per well. The cells were then restimulated with 10 μg/ml of MOG_p35-55_ and incubated at 37°C for 5 days. Concanavalin A (Con A) at a concentration of 2 mg/ml and media was used as controls. On day 6 the cells were collected, blocked with anti-Fc block antibody, stained with APC CD4 antibody and acquired and analyzed by flow cytometry.

### RNA sequencing and analysis

The study cohort and procedures were previously described ^11^. Briefly, 11 subjects with inflammatory demyelinating disease (9 with RRMS and 2 with anti-MOG disorder) and 3 control subjects (1 with amyotrophic lateral sclerosis [ALS], 1 with idiopathic intracranial hypertension [IIH], and 1 healthy control [HC]) were recruited for a study to assess the characteristics of CSF and blood cells.

On the day of cell collection using fresh cells, droplet-based 39 (for anti-MOG disorder subject 1) and 59 (for subjects 2 and 5 with RRMS) libraries were prepared using Chromium Single Cell 39 v2 or 59 Reagent Kits according to the manufacturer’s protocol from 10× Genomics. The generated scRNA libraries were sequenced using an Illumina HiSeq4000 or Novaseq sequencer. Cell Ranger Single-Cell Software 2.2 was used to perform sample demultiplexing, barcode processing, and single-cell 39 and 59 counting. Afterward, fastqfiles for each sample were processed with a cellranger count, which was used to align the samples to GRCh38 genome, filter, and quantify reads. For each sample, the recovered-cells parameter was specified as 10,000 cells that we expected to recover for each individual library. To reanalyze scRNA-seq data performed with SeqWell publicly available from GSE117397, we downloaded count tables from 2 subjects with HIV for whom both CSF and blood samples were available from GEO DataSets (GSM3293822, HIV1_Bld; GSM3293823, HIV1_CSF; GSM3293824, HIV2_Bld; GSM3293825, HIV2_CSF). To combine data from all samples together, we used the canonical correlation analysis (CCA) algorithm from Seurat 3 ^12^. Before applying CCA, we removed cells from the 39 and 59 data sets that contain more than 10% of mitochondrial RNA. Cells from SeqWell data sets with less than 20% of RNA from mitochondrial genes and more than 500 but less than 2,500 expressed genes were considered viable cells. After CCA data were scaled, principal component analysis (PCA) was performed with RunPCA function. A Uniform Manifold Approximation and Projection (UMAP) dimensionality reduction was performed on the scaled matrix using the first 20 PCA components to obtain a 2-dimensional representation of the cell states ^13^. For clustering, we used FindNeighbours and FindClusters functions on 20 PCA components with a resolution of 0.5. FindAllMarkers function was used to characterize clusters. For heatmap representation, the mean expression of markers inside each cluster was used. Heatmaps were built with Phantasus software (artyomovlab.wustl.edu/phantasus/). Applying this method, data from a total of 23,363 PBMCs and 14,179 CSF cells were combined and visualized using the UMAP technique 15 ^13^ and unsupervised clustering ^14,15^.

### Statistical analysis

Groups were compared for normality using the Kolmogorov-Smirnov test. Normally distributed data were compared using unpaired two-sided Student *t* test. Holm-Sidak post hoc analysis was performed to correct in case of multiple comparisons. All statistical analyses were 2-sided and a P value of less than 0.05 was set as the level of statistical significance. In the figures, P values are shown as follows: * = 0.05; ** = 0.005; *** = 0.0005; **** = 0.00005; ***** = 0.000005. Each experiment was repeated at least once. All analyses were performed with Prism 8 (Graphpad, La Jolla, CA).

## Results

### Reduction of α4-integrin expression in naïve CD11c.Cre^+/-^*ITGA4*^fl/fl^ mice

We first assessed the effectiveness of Cre-Lox recombination on the expression of α4-integrin on the surface of CD11c^+^ leukocytes in CD11c.Cre^+/-^*ITGA4*^fl/fl^ mice. The percentage of α4-integrin expressing CD11c^+^ leukocytes, collected from primary and secondary lymphoid compartments of naïve WT and CD11c.Cre^+/-^*ITGA4*^fl/fl^ mice was assessed using *ex vivo* flow cytometry. When compared with naïve WT controls, naïve CD11c.Cre^+/-^*ITGA4*^fl/fl^ mice had a significantly lower frequency of CD11c^+^ cells expressing α4-integrin in spleen (Figure 1A), lymph nodes (Figure 1B), bone marrow (Figure 1C) and blood (Figure 1D) (P value < 0.0005). This observation confirms that Cre-mediated gene recombination diminished the expression of α4-integrin in CD11c.Cre^+/-^*ITGA4*^fl/fl^ mice. We also confirmed a similar α4-integrin expression by CD11c^+^ leukocytes between WT and *ITGA4*^fl/fl^ mice across multiple generations, which were found to be indistinguishable (data not shown). In subsequent experiments, we only compared results between CD11c.Cre^+/-^*ITGA4*^fl/fl^ mice and WT controls.

**Figure 1.**
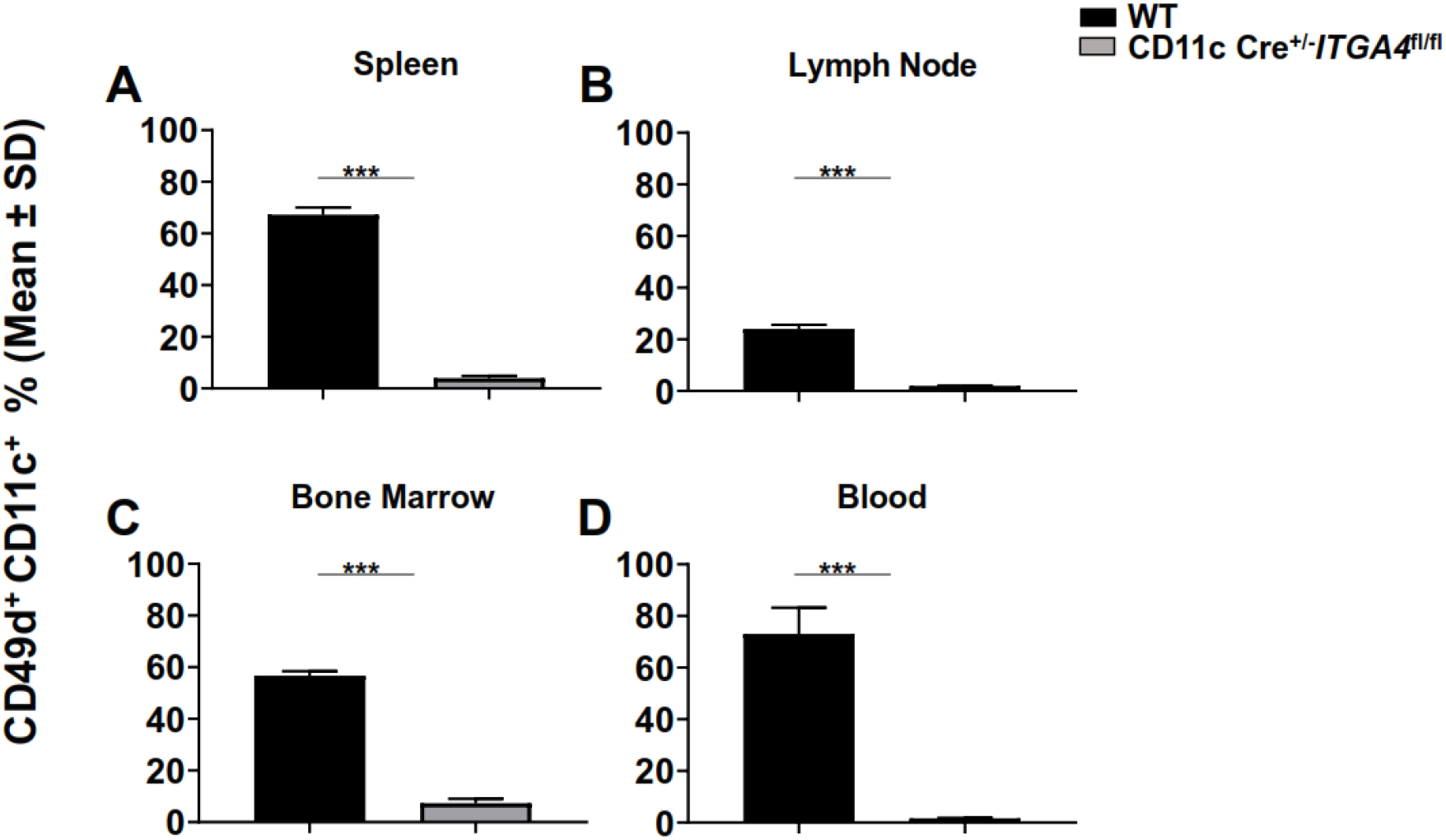
CD11c.Cre^+/-^*ITGA4*^fl/fl^ mice exhibited reduced expression of α4-integrin by CD11c^+^ cells. The mean ± standard deviation (SD) of α4-integrin (CD49d)-expressing CD11c^+^ cells (%) from total 50 x 10^3^ recorded viable cells in (**A**) spleen, (**B**) lymph node, (**C**) bone marrow, and (**D**) blood of naïve wild type and CD11c.Cre^+/-^*ITGA4*^fl/fl^ mice is shown (N = 9 experimental animals per group; data show the pooled analysis of all study cohorts). The α4-integrin expression in CD11c.Cre^+/-^*ITGA4*^fl/fl^ mice compared to C57BL/6 wild type (WT) controls is significantly reduced in all compartments (P value < 0.0005).

### Clinical onset of active EAE is delayed in CD11c.Cre^+/-^*ITGA4*^fl/fl^ mice

Based on our observations in a post-autopsy study of PML related to natalizumab therapy ^4^ and the critical role of DC in the initiation and perpetuation of CNS autoimmunity, we hypothesized that limited expression of α4-integrin by CD11c^+^ cells would lead to reduction of severity of clinical EAE scores. Active EAE was induced in CD11c.Cre^+/-^*ITGA4*^fl/fl^ mice and WT controls by immunization with MOG_p35-55_ in CFA. There was no significant difference between CD11c.Cre^+/-^*ITGA4*^fl/fl^ mice and WT controls regarding disease severity following active immunization (**Figure 2A**). However, the onset of clinical EAE was significantly and reproducibly delayed by at least one day in CD11c.Cre^+/-^*ITGA4*^fl/fl^ mice as compared with WT controls (**Figure 2B**). Going forward, we will refer to the delay in disease onset as “EAE onset lag”. Moreover, we will refer to an EAE phenotype that continues beyond a brief monophasic clinical episode as “persistent clinical EAE”. The EAE onset lag allowed us to characterize cellular events in relevant tissues during the preclinical stage and the early onset of EAE in CD11c.Cre^+/-^*ITGA4*^fl/fl^ mice in comparison to WT controls and characterize the contribution of various cellular immune populations. *Ex vivo* flow cytometry analyses of the peripheral lymphoid tissues and CNS compartment during acute and ongoing EAE stages were performed.

**Figure 2.**
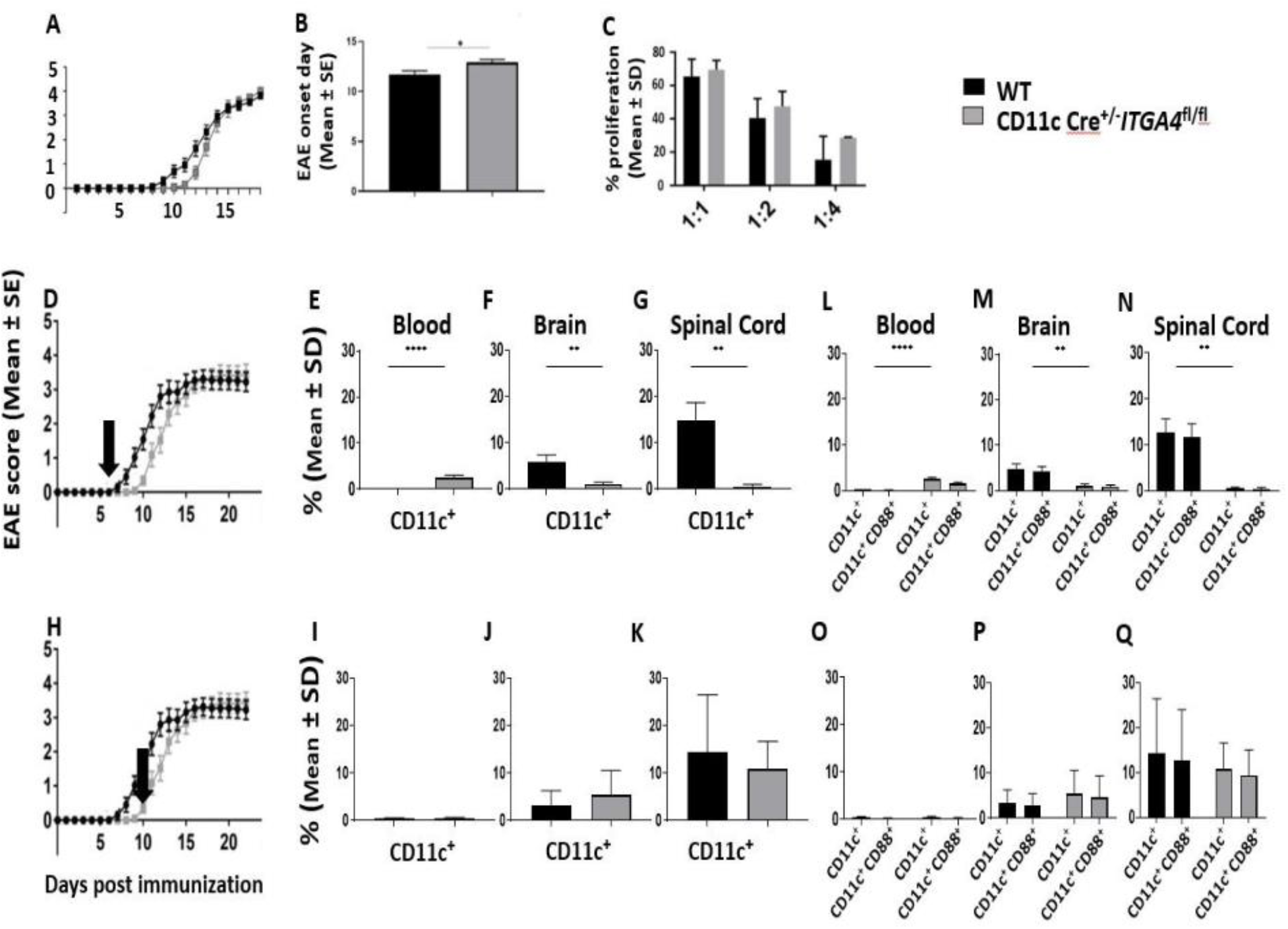
The onset of active experimental autoimmune encephalomyelitis (EAE) is consistently delayed among the CD11c.Cre^+/-^*ITGA4*^fl/fl^ mice. (**A**) The mean ± standard error (SE) of clinical EAE following active immunization are presented (N= 24 experimental animals per group; data display pooled analysis of all study cohorts). (**A & B**) The onset of disease was significantly delayed in the CD11c.Cre^+/-^*ITGA4*^fl/fl^ group (P value < 0.05). (**C**) The percentage (mean ± standard deviation (SD)) of proliferating Ag-specific CD4^+^ T cells co-cultured with different dilutions of CD11c^+^ splenocytes from CD11c.Cre^+/-^*ITGA4*^fl/fl^ mice and WT controls is shown. There was no significant difference between the two groups. Clinical EAE is preceded by accumulation of CD11c^+^ cells in peripheral blood. The mean ± standard deviation (SD) of percentage of CD11c^+^ and CD11c^+^CD88^+^ cells from total 50 x 10^3^ recorded viable cells upon the onset of actively-induced clinical EAE in (**D**) C57BL/6 wild type (WT) controls, in (**E, L**) blood, (**F, M**) brain, and (**G, N**) spinal cord, and after the onset of clinical EAE in CD11c.Cre^+/-^*ITGA4*^fl/fl^ mice (**H**) in (**I, O**) blood, (**J, P**) brain, and (**K, Q**) spinal cord and is shown (N = 8 experimental animals per group; data show pooled analysis of all study cohorts). (**E, I**) There was a significant sequestration of CD11c^+^ and CD11c^+^CD88^+^cells in the blood of CD11c.Cre^+/-^*ITGA4^fl/fl^* compared to controls upon the onset of clinical EAE (P value < 0.00005), while CD11c^+^ cells were significantly more frequent in the (**F, J**) brain and (**G, K**) spinal cord of wild type mice (P value 0.005). (**F, G, H**) There was no difference between the two groups in any of the compartments after the start of the clinical EAE among the CD11c.Cre^+/-^*ITGA4*^fl/fl^ mice (P value > 0.05).

### Antigen presentation is intact in CD11c.Cre^+/-^*ITGA4*^fl/fl^ mice

Previous studies indicated that blocking very late-activating antigen 4 (VLA-4; α4β1-integrin) with mAb negatively impacted the ability of CD4^+^ T cells to proliferate, perhaps suggesting a role for α4-integrin as a co-stimulatory molecule ^16,17^. We assessed this in CD11c.Cre^+/-^*ITGA4*^fl/fl^ mice and WT controls through flow cytometry based recall proliferation assays. MOGp35-55-specific CD4^+^ T cells were co cultured with different ratios of CD11c^+^ APC harvested from CD11c.Cre^+/-^*ITGA4*^fl/fl^ mice and WT controls. *Ex vivo* flow cytometry assessment of the CD4^+^ T cells proliferation based on CFSE dilution indicated that there was no significant difference between CD11c.Cre^+/-^*ITGA4*^fl/fl^ mice and WT controls regarding the recall response by the CD4^+^ T cells (**Figure 2C**). This suggested that antigen (Ag) presentation and T cell activation remained intact in CD11c.Cre^+/-^*ITGA4*^fl/fl^ mice, despite a significantly lower expression of α4-integrin by APC.

### Leukocyte composition aside from the CD11c cell lineage is unaltered in CD11c.Cre^+/-^*ITGA4*^fl/fl^ mice

Next, we investigated whether the delayed onset of clinical EAE in CD11c.Cre^+/-^ *ITGA4*^fl/fl^ mice was accompanied by an altered composition of leukocytes in different compartments. Flow cytometry analysis of immune cell populations revealed that leukocyte subtypes including CD45^+^, CD19^+^, CD4^+^ and CD8^+^ cells, from CD11c.Cre^+/-^*ITGA4*^fl/fl^ mice and WT controls had similar composition in different compartments during naïve states as well as during acute and persistent clinical EAE (**Supplementary Figure 2**). These observations suggest that the observed EAE onset lag was not driven by impaired access of immune cell outside the CD11c^+^ lineage to the CNS compartment.

### CD11c^+^ cells sequester in peripheral blood prior to the onset of clinical active EAE

As shown earlier, we observed that the onset of clinical EAE following active immunization was consistently delayed in the CD11c.Cre^+/-^*ITGA4*^fl/fl^ group. The EAE onset lag provided a window for studying the composition of the inflammatory milieu in different compartments. Using *ex vivo* flow cytometry, we compared the percentage of live CD45^+^MHCII^+^CD11c^+^ cells collected from the peripheral blood and CNS (brain and spinal cord) of CD11c.Cre^+/-^*ITGA4*^fl/fl^ mice and WT controls at two time points following induction of active EAE. The cells were collected from both groups upon the onset of clinical EAE among the WT controls (**Figure 2D**). In this setting, CD11c^+^ cells were significantly more frequent in the blood of CD11c.Cre^+/-^*ITGA4*^fl/fl^ mice compared with WT controls (**Figure 2E**) (P value < 0.00005); simultaneously, these cells were significantly more abundant in the brain (**Figure 2F**) and spinal cord (**Figure 2G**) of the WT controls (P value < 0.005). Subsequently, cells were collected from mice in both groups at the onset of clinical EAE among the CD11c.Cre^+/-^*ITGA4*^fl/fl^ mice (**Figure 2H**). There were no significant differences between the frequencies of CD11c^+^ cells inside the peripheral circulation (**Figure 2I**), brain (**Figure 2J**), and spinal cord (**Figure 2K**) of the CD11c.Cre^+/-^*ITGA4*^fl/fl^ mice compared to WT controls. Taken together, these findings indicated that the clinical onset of EAE was closely related to the increased appearance of CD11c^+^ cells inside the CNS compartment. The concomitant disappearance of these cells from the blood suggested that CD11c^+^ cells inside the CNS originated from cells entering from the peripheral compartment.

### CD11c^+^ cells and CD11c^+^CD88^+^ cells distribute similarly among tissues during active EAE

We observed that the onset of clinical EAE was preceded by the accumulation of CD11c^+^ cells in the blood, and their subsequent appearance inside the CNS was associated with the onset of clinical disease. Next, we wanted to further characterize immunologic traits of these cells. Bone marrow-derived CD11c^+^ myeloid cells in the blood are comprised to a significant extent of CD88^+^ monocyte-derived cells ^18–20^. Using *ex vivo* flow cytometry, we assessed the percentage of CD88 expressing CD11c^+^ cells collected from the blood and CNS of CD11c.Cre^+/-^*ITGA4*^fl/fl^ mice and WT controls. In both groups, the majority of CD11c^+^ cells in the blood and CNS also expressed CD88. At the time of the onset of clinical EAE in the WT control group (**Figure 2D**), CD11c^+^ CD88^+^ cells accumulated in the blood of CD11c.Cre^+/-^*ITGA4*^fl/fl^ mice (**Figure 2L**) (P value < 0.00005), while they were significantly more frequent inside the brain (**Figure 2M**) and spinal cord (**Figure 2N**) of WT controls (P value < 0,005). Subsequently, following the onset of clinical EAE in CD1c.Cre^=/-^*ITGA4*^fl/fl^ mice (**Figure 2H**), the percentage of CD11c^+^ CD88^+^ cells in the blood was diminished (**Figure 2O**), while it was increased in the brain (**Figure 2P**) and spinal cord (**Figure 2Q**). At that time, there were no differences between the two experimental groups in any of these compartments.

### CD11c^+^CD88^+^ cells express CD317 inside the CNS but not in blood

We observed that CD11c^+^ cells from CD11c.Cre^+/-^*ITGA4*^fl/fl^ mice were initially delayed from gaining access to the CNS, but ultimately accumulated inside the brain and spinal cord of CD11c.Cre^+/-^*ITGA4*^fl/fl^ mice (**Figure 2**). This suggested redundancy in adhesion molecules by CD11c^+^ myeloid cells. CD317 is a lipid raft associated glycosyl-phosphatidylinositol (gpi)-anchored protein that is constitutively expressed by plasmacytoid DC, mature B cells, plasma cells, and other leukocytes ^21,22^. More recently, CD317 has also been shown to function as an adhesion molecule for myeloid cells ^23^. To determine whether CD317 is upregulated by CD11c^+^CD88^+^ cells upon entry into the CNS and serves as an adhesion molecule, we assessed its expression by CD11c^+^ cells in our model. During active EAE, we found that CD317 expressing CD11c^+^ CD88^+^ cells in the peripheral circulation were significantly less frequent (P value < 0.0005) than CD11c^+^CD88^+^CD317^-^ cells (**Figure 3A**). In contrast, the percentage of CD11c^+^CD88^+^ cells expressing CD317 was significantly higher than the percentage of CD11c^+^ CD88^+^ cells not expressing CD317 in the brain (**Figure 3B**) and spinal cord (**Figure 3C**) (P value <0.0005). Together, these observations suggest that CD11c^+^CD88^+^ cells in the peripheral circulation start expressing CD317 upon entry into the CNS compartment at the onset of clinical EAE.

**Figure 3.**
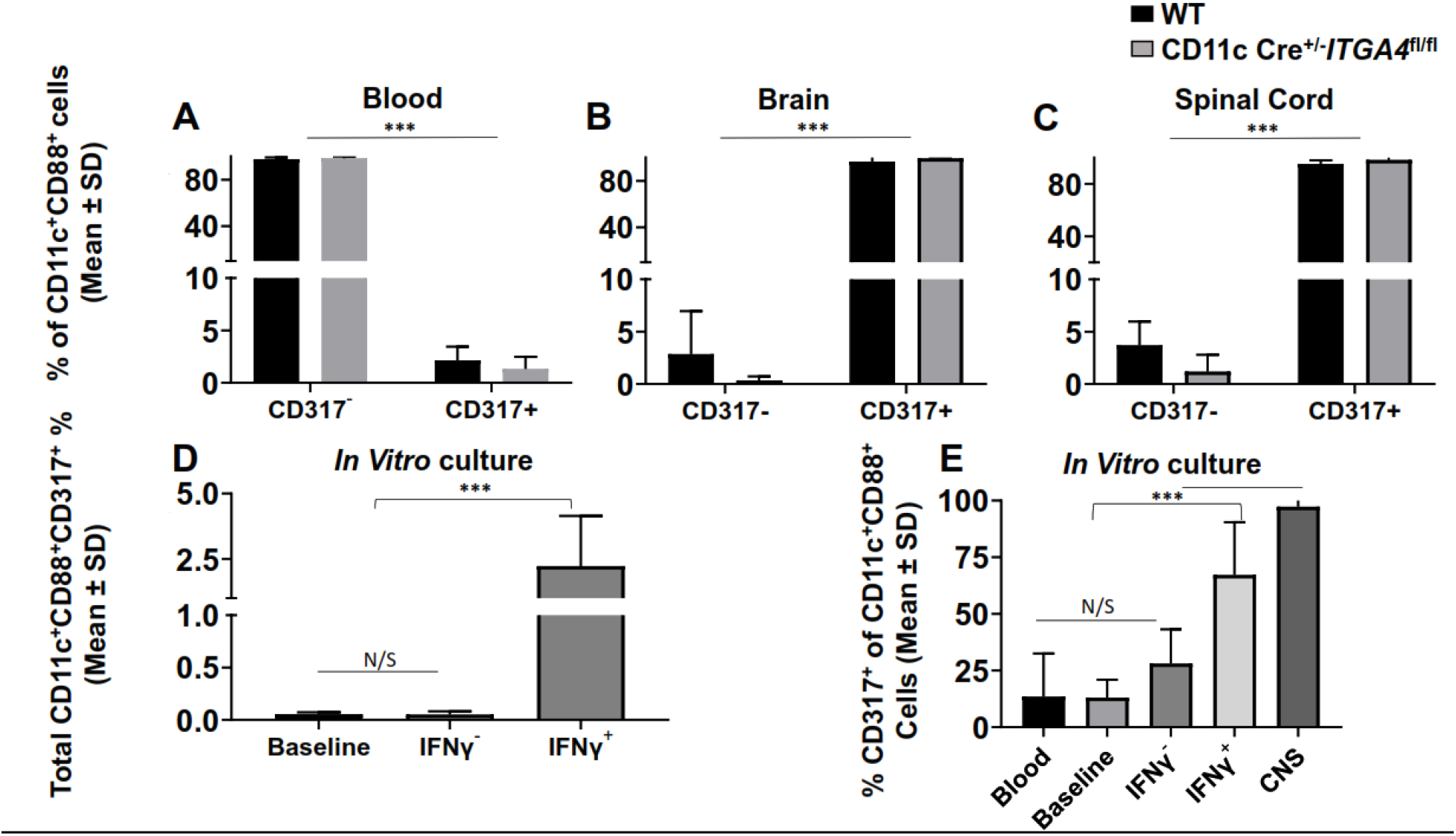
CD11c^+^CD88^+^ cells start expressing CD317 inside the CNS compartment at the onset of active clinical experimental autoimmune encephalomyelitis (EAE). The mean ± standard deviation (SD) of the percentage of CD11c^+^ CD88^+^ cells that express CD317 in (**A**) blood, (**B**) brain, and (**C**) spinal cord during actively-induced EAE in CD11c.Cre^+/-^*ITGA4*^fl/fl^ mice or C57BL/6 wild type (WT) controls is shown (N = 8 experimental animals per group; data show pooled analysis of all study cohorts). CD11c^+^CD88^+^ cells in the (**A**) blood are mostly CD317^-^, while inside the (**B, C**) brain and spinal cord CD317^+^ cells constitute the majority of the CD11c^+^CD88^+^ cells (P value < 0.0005). (**D**) The mean ± SD of CD11c^+^ CD88^+^ CD317^+^ cells following *in vitro* culture (N =12 replicates per group; data show pooled analysis of all study cohorts). (**D**) Interferon gamma (IFN γ) significantly upregulates the expression of CD317 in CD11c^+^CD88^+^ cells (P value < 0.0005). (**E**) The mean ± SD of percent of CD317^+^ cells from the total CD11c^+^CD88^+^ cells is shown. The level of IFNγ-induced expression of CD317 in CD11c^+^CD88^+^ cells resembles that of the expression levels of CD317 in CD11c^+^CD88^+^ cells recovered from the CNS of mice actively immunized for EAE. Their frequency is significantly higher than baseline cells, cells recovered from blood of actively immunized mice and cultured cells that did not receive IFN γ treatment (P value < 0.0005)

The expression of CD317 is inducible through engagement of Signal Transducer And Activator Of Transcription (STAT) following exposure to type I or type II IFN ^24–26^ To investigate whether CNS CD11c^+^CD88^+^CD317^+^ cells originate from blood CD11c^+^CD88^+^ cell at the onset of clinical EAE, we collected peripheral blood mononuclear cells (PBMC) from WT mice immunized with MOG_p35-55_, prior to the expected onset of clinical EAE. *In vitro* culture was performed with or without stimulation with interferon gamma (IFNγ). Subsequent flow cytometry analysis of the cultured cells revealed that CD11c^+^CD88^+^ cells that were exposed to IFNγ in culture upregulated CD317 on their cell surface (**Figure 3D**). The percentage of CD11c^+^CD88^+^ CD317^+^ was significantly higher in the IFNγ-treated group (P value < 0.0005). Interestingly, the fraction of CD11c^+^CD88^+^ cells expressing CD317 after *in vitro* exposure to IFNγ was comparable to that of these cells observed in the CNS compartment during clinical EAE in both experimental groups, and significantly higher than that of blood CD11c^+^CD88^+^ cells (**Figure 3E**).

### CD11c^+^CD88^+^ cells and CD11c^+^CD88^+^CD317^+^ cells express CD11b

To characterize blood CD11c^+^CD88^+^ cells during preclinical EAE as well as CNS CD11c^+^CD88^+^CD317^+^ cells during established EAE, we assessed both cell types for CD11b expression. CD11 b is expressed on granulocyte, macrophages, monocytes, and natural killer cells ^27^. We observed that CD11c^+^CD88^+^ cells and CD11c^+^CD88^+^CD317^+^ cells also express CD11b (**Supplementary Figure 1**), a property that has been assigned to migratory DC ^27^.

### Decreased EAE severity in the absence of CD11c^+^CD88^+^CD317^+^ cells in the CNS

In active EAE, the microenvironment outside of the CNS in this EAE model is defined by an excess of IFNγ ^28^. In contrast, the adoptive transfer of myelin-reactive CD4^+^ T cells, many of which are Th1 cells, creates an IFNγ-rich environment inside the CNS, but not in other tissues ^29–31^. MOG_p35-55_-reactive CD4^+^ T cells from WT donors were transferred to CD11c.Cre^+/-^*ITGA4*^fl/fl^ and WT controls recipients (**Figure 4A**). Analysis of clinical disease scores in both groups longitudinally revealed a striking reduction of clinical disease in the CD11c.Cre^+/-^*ITGA4*^fl/fl^ mice as compared with WT controls (P value < 0.0005) (**Figure 4A**). Specifically, the disease course was substantially ameliorated and abbreviated, and devoid of the persistent clinical phase that is routinely expected in the C57BL/6 model of EAE. *Ex vivo* flow cytometry revealed that CD11c^+^CD88^+^ cells were significantly more frequent in the blood of CD11c.Cre^+/-^ *ITGA4*^fl/fl^ mice compared to WT controls at day 18 post adoptive transfer (**Figure 4B**) (P value < 0.005). Further *ex vivo* flow cytometry-based characterization of immune cells in the peripheral compartment and CNS of mice in both groups indicated that CD11c.Cre^+/-^*ITGA4*^fl/fl^ mice that showed less severe disease course also exhibited significantly lower percentage of CD11c^+^ cells, CD11c^+^CD88^+^ cells, and CD11c^+^CD88^+^CD317^+^ cells inside the brain (**Figure 4C,F**) and spinal cord (**Figure 4D,G**) (P value < 0.005). Interestingly, CD11c^+^CD88^+^CD317^+^ were extremely rare in the blood of both CD11c.Cre^+/-^*ITGA4*^fl/fl^ mice and WT controls (**Figure 4E**). In summary, these observations suggest an association between the presence of CD11c^+^CD88^+^CD317^+^ cells inside the CNS and the severity of acute EAE, as well as the transition to persistent clinical EAE.

**Figure 4.**
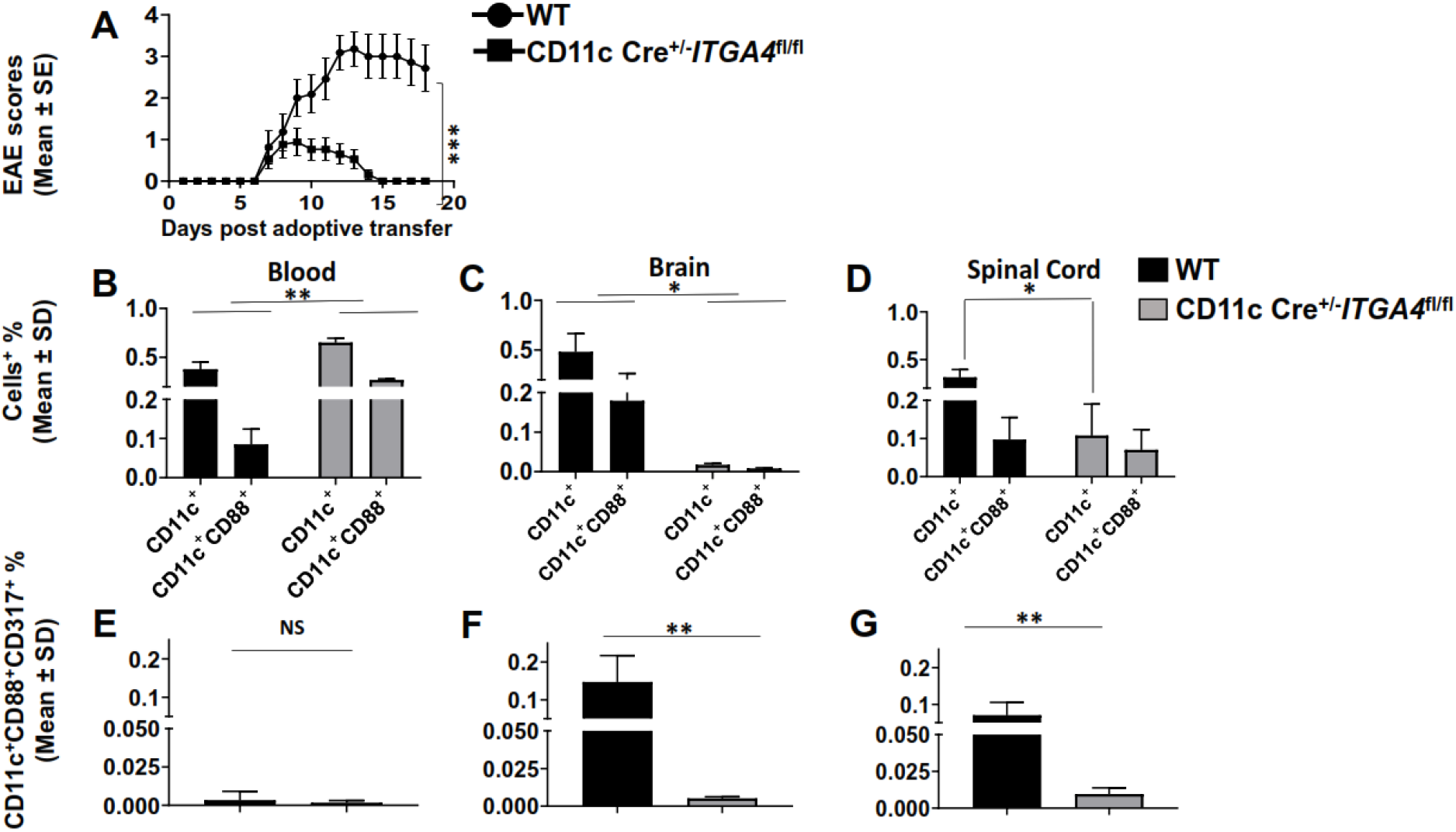
Adoptive transfer of MOGp35-55-specific CD4^+^ T cells into CD11c.Cre^+/-^*ITGA4*^fl/fl^ mice causes ameliorated experimental autoimmune encephalomyelitis (EAE). (**A**) Adoptive transfer EAE was induced by transferring myelin oligodendrocyte glycoprotein (MOG) peptide (p) 35-55 (MOGp35-55)-reactive CD4^+^ T cells from C57BL/6 wild type (WT) mice into CD11c.Cre^+/-^ *ITGA4*^fl/fl^ mice or C57BL/6 wild type (WT) controls. The mean ± standard error (SE) of clinical EAE scores following adoptive transfer are presented (N = 11 experimental animals per group; data show pooled analysis of all study cohorts). Clinical disease among the CD11c.Cre^+/-^*ITGA4*^fl/fl^ was significantly ameliorated (P value < 0.0005). The percentages of CD11c^+^ cells, CD11c^+^CD88^+^ cells (mean ± SD) out of 50 x 10^3^ recorded viable cells in (**B**) blood, (**C**) brain, and (**D**) and spinal cord during clinically persistent EAE in the WT are shown (N = 3 experimental animals per group; data show pooled analysis of all study cohorts). The percentage of CD11c^+^CD88^+^CD317^+^ cells (mean ± SD) out of 50 x 10^3^ recorded viable cells is also shown in (**E**) blood, (**F**) brain, and (**G**) spinal cord. There was a significant reduction of the (**C, D**) CD11c^+^CD88^+^ and (**F, G**) CD317^+^ cells in the CNS of CD11c.Cre^+/-^*ITGA4*^fl/fl^ compared to WT controls (P value < 0.05). CD11c^+^ CD88^+^ cells were significantly more frequent in the (**A**) blood of CD11c.Cre^+/-^*ITGA4*^fl/fl^ mice compared to WT controls (P value < 0.005) (N = 3 experimental animals per group; data show pooled analysis of all study cohorts).

### Clustering of CD11c^+^CD88^+^CD317^+^cells within microglia-like cells in the CNS and CSF by transcriptomic profiling

After identifying the association of CD11c^+^CD88^+^CD317^+^ cells with EAE disease severity, we aimed to determine whether this cell type could also be identified in human tissues relevant to CNS autoimmunity. Using scRNA-seq, our group previously described 20 clusters representing distinct immune cell populations within PBMC and CSF of subjects with neurologic diseases ^11^. Characteristic gene expression markers were employed to assign cellular identities to each cluster of myeloid populations in blood (**Figure 5A**) and CSF (**Figure 5B**). Recent studies identified myeloid cells within the CSF using scRNA-seq that resembled microglia ^11^,^32–34^, which are characterized by the expression of signature genes including CX3CR1, CSF1R, SLC2A5, MARCKS, and P2RY13 ^35–37^.

**Figure 5.**
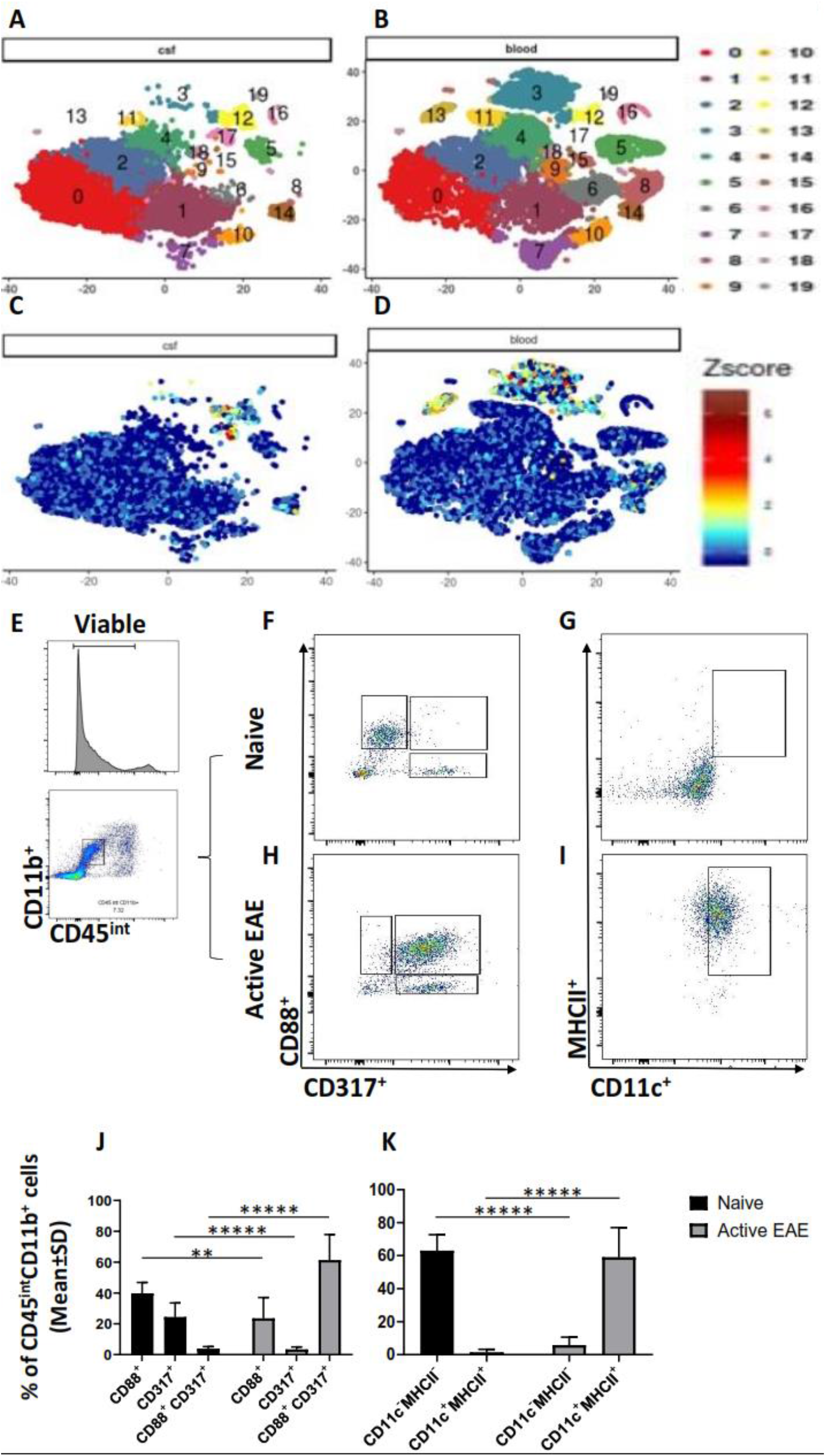
CD11c^+^CD88^+^CD317^+^cells localize within microglia cells in the CNS and CSF. Using scRNA-seq, our group previously described 20 clusters representing distinct cell populations were identified in PBMC and CSF ^11^. Characteristic gene expression markers were employed to assign cellular identities to each cluster of myeloid populations in (**A**) CSF, and (**B**) blood. Three prior studies identified myeloid CSF cells using scRNA-seq that resembled microglia in cluster 17, and CD11c^+^CD88^+^CD317^+^cells localize within this cell population. Activated, but not naive microglia express CD11c, CD88, and CD317. Active experimental autoimmune encephalomyelitis (EAE) was induced in C57BL/6 wild-type (WT) mice. Brains and spinal cords were isolated and dissociated. (**E**) Microglia were defined as viable CD45 intermediate (int), CD11b^+^ cells. (**F**) In microglia from naïve cells, these cells did not express CD88 and CD317 concomitantly. (**G**) Moreover, naïve microglia are not high expressers of MHC II and CD11c. In contrast, microglia from mice with established active EAE express high levels of (**H**) CD88 and CD317, as well as (**I**) MHC II and CD11c. The differences in the expression of (**J**) CD88, CD317, MHC II, and (**K**) CD11c were highly significant (N = 6 experimental animals per group; data show pooled analysis of all study cohorts; (P value < 0.005 or < 0.000005).

The expression profile for *ITGAX* (CD11c), *C5AR1* (CD88), and *BST2* (CD317) within CSF myeloid cell populations revealed a preponderance within the cluster of cells designated as microglia (**Figure 5C**). These cells were undetectable in blood (**Figure 5D**). This finding indicates that CD11c^+^CD88^+^CD317^+^cells are CNS compartment specific and share a transcriptional profile with myeloid cells previously identified as microglia.

### Activated, but not naïve, microglia express CD11c, CD88, and CD317

Intrigued by the finding that human CSF CD11c^+^CD88^+^CD317^+^ cells localize within microglia cells by transcriptomic analyses, we aimed to determine whether parenchymal microglia also express these surface markers. Due to the lack of human CNS tissue, we decided to assess murine microglia cells. Microglia were defined as viable CD45 intermediate (int), CD11b^+^ cells (**Figure 5E**). Microglia from naïve mice did not express CD88 and CD317 concomitantly (**Figure 5F**). Moreover, naïve microglia are not high expressers of MHC II and CD11c (**Figure 5G**). In contrast, microglia from mice with established active EAE express high levels of CD88 and CD317 (**Figure 5H, J**), as well as MHC II and CD11c (**Figure 5I, K**). In summary, these data indicate that CNS CD11c^+^CD88^+^CD317^+^ cells associated with CNS autoimmune disease resemble activated parenchymal microglia.

### Anti-CD317 treatment ameliorates EAE and shortens disease duration

To provide further evidence that CD11c^+^CD88^+^CD317^+^ cells are critical for establishment of persistent clinical EAE, and are therefore potential therapeutic targets, we administered a commercially available anti-CD317 rat-anti-mouse IgG2b mAb on days 7 and 8 post active immunization (**Figure 6A**). Anti-CD317 treatment ameliorated acute EAE, with complete resolution of clinical disease. Anti-CD317 therapy resulted in an increase of CD11c^+^, CD11c^+^CD88^+^, and CD11c^+^CD88^+^CD317^+^cells in blood (**Figure 6B**). However, the difference in cell numbers compared to vehicle controls group was not statistically significant. Furthermore, in the treatment group, CD11c^+^, CD11c^+^CD88^+^, and CD11c^+^CD88^+^CD317^+^cells in the brain (**Figure 6C**) and spinal cord (**Figure 6D**) were reduced compared with controls. This was statistically significant in the spinal cord (**Figure 6D**) (P value < 0.005). Differences in the percentages of CD11c^+^ cells and CD11c^+^CD88^+^ in the blood, brain, and spinal cord in the anti-CD317 treatment group were driven by changes in the percentages of CD11c^+^CD88^+^CD317^+^cells. Our observations suggest that anti-CD317 therapy in EAE diminishes the ability of CD11c^+^CD88^+^CD317^+^ cells to gain access to the CNS, reducing the severity of acute EAE and the transition to persistent clinical EAE.

**Figure 6.**
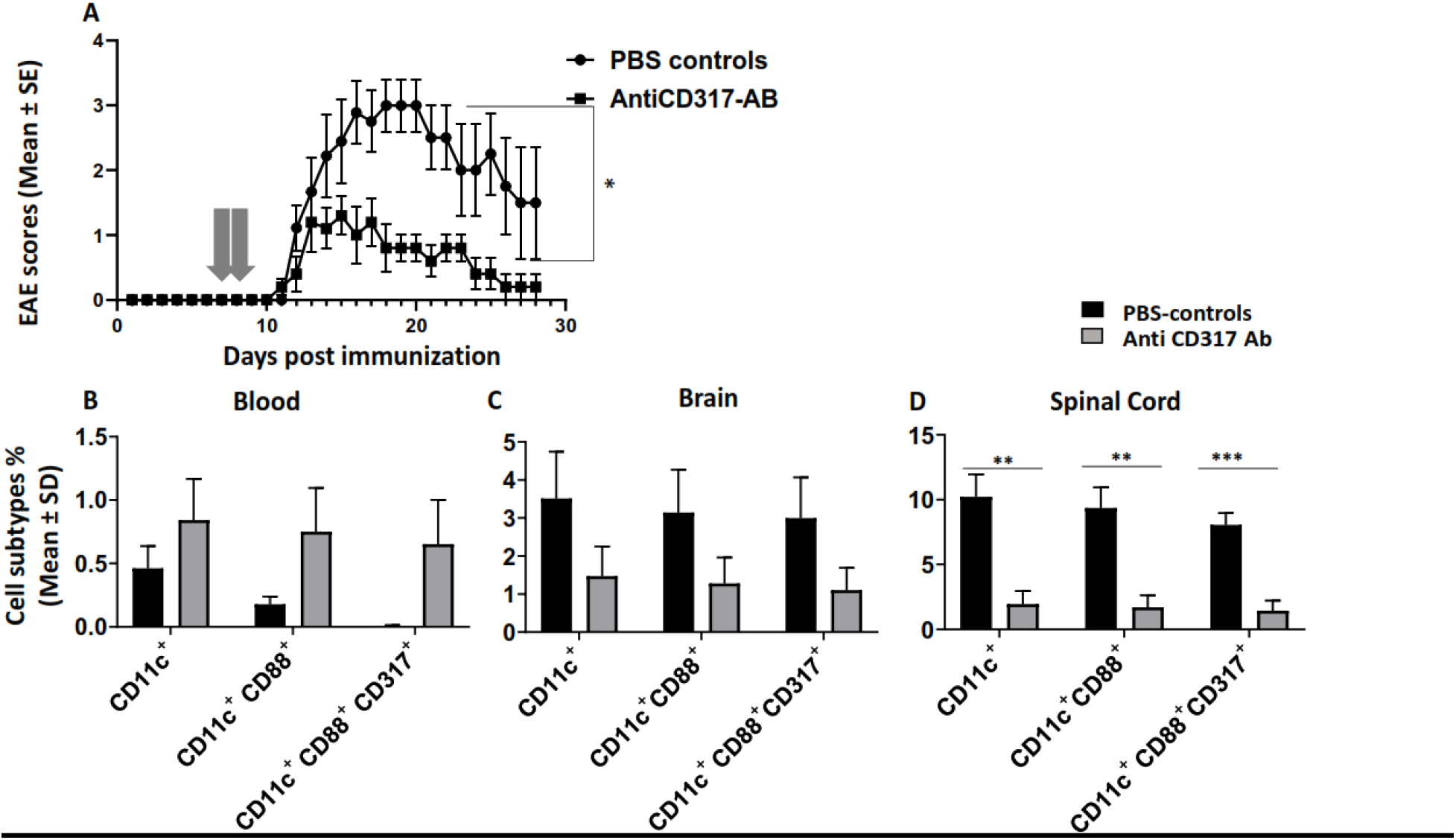
Anti-CD317 treatment prevents established experimental autoimmune encephalomyelitis (EAE). (**A**) EAE was actively induced in C57BL/6 wild-type (WT) mice. The mean ± standard error (SE) of clinical EAE scores following the injection of anti-CD317 monoclonal antibody (mAb) in active EAE are shown (N =10 experimental animals per group; data show pooled analysis from all study cohorts). There was a significant amelioration of clinical scores in the anti-CD317 mAb treatment group compared to PBS controls (P value < 0.05). (**B**) The mice in the anti-CD317 mAb treatment group showed a trend of sequestration of CD11c^+^ cells, CD11c^+^CD88^+^ cells, and CD11c^+^CD88^+^CD317^+^ cells in the blood however the differences were not statistically significant (N = 4 experimental animals per group; data show pooled analysis of all study cohorts). In the anti-CD317 mAb treatment group, CD11c^+^ cells, CD11c^+^CD88^+^ cells, and CD11c^+^CD88^+^CD317^+^ cells were less frequent in the (**C**) brain and (**D**) spinal cord. A statistically significant difference was observed in the spinal cords between the two treatment groups (P value < 0.005 and 0.0005).

## Discussion

In the present study, we investigated the effects of selective α4-integrin deletion in CD11c^+^ cells in the EAE model of MS. Our observations indicate that the appearance of CD11c^+^CD88^+^CD317^+^ myeloid cells inside the CNS compartment is required for establishment of persistent clinical EAE after active immunization. This is further supported by the fact that reducing the ability of these cells to gain access to the CNS via a blocking anti-CD317 mAb also prevented persistent clinical EAE. This suggests CD317 as a promising molecular target for treatment in patients with MS and related disorders.

Classically, CD11c^+^ myeloid cells are believed to participate in processing and presentation of antigens to Ag-specific T cells during CNS autoimmunity in secondary lymphoid organs and the brain and spinal cord ^2^. Based on our results, we suggest that the role of CD11c^+^CD88^+^CD317^+^ cells in EAE pathogenesis may be largely independent of antigen presentation. CD88^+^ DC were previously shown to have minimal antigen presentation capability during active EAE as evidenced by a limited ability to induce T cell proliferation to recall antigens ^20^. In fact, initiation of clinical EAE, which requires Ag processing and presentation in peripheral lymphatic tissues, is mainly driven by classical tissue-resident cDC, characterized by their expression of CD26 ^20^. We showed in our model that Ag presentation and ability to induce T cell proliferation was maintained despite the deletion of α4-integrin in CD11c cells in Cre^+/-^*ITGA4*^fl/fl^ mice. Furthermore, our results indicated that the pathogenic role of CD11c^+^CD88^+^CD317^+^ cells in EAE appeared dependent on their physical presence in the inflammatory environment in the CNS, since inhibiting their entry into the CNS compartment with anti-CD317 mAb ameliorated clinically EAE, but did not eliminate these cells from the peripheral blood.

CD11c^+^ cells from CD11c.Cre^+/-^*ITGA4*^fl/fl^ mice have a significantly reduced expression of α4-integrin. In spite of this observation, CD11c^+^ cells are merely delayed in, but not prevented from gaining access to the CNS. Ultimately CD11c^+^ cells are abundantly present inside the brain and spinal cord of CD11c.Cre^+/-^*ITGA4*^fl/fl^ mice at EAE disease onset following active immunization. We posit that this intriguing observation may point to a redundancy in the usage of adhesion molecules in CD11c^+^ myeloid cells in gaining access to the CNS compartment. Work by other investigators suggests that macrophages utilize CD317 as an adhesion molecule to gain access to their target tissue during inflammation ^23^. In our study, CD11c^+^CD88^+^ cells circulated and sequestered in the peripheral blood of CD11c.Cre^+/-^*ITGA4*^fl/fl^ mice, and it was only after they acquired the expression of CD317 that they gained access to the CNS. The exact role and mechanism how CD317 expression endows these cells with access to the CNS, and whether it serves directly as an alternative adhesion molecule that substitutes or complements α4-integrin, needs further studies.

As mentioned earlier, CD317 is constitutively expressed by many cell types, including myeloid cells and B lymphocytes ^21,22^. There are species-specific differences regarding the expression of CD317. While human plasmacytoid DC express low amounts of surface CD317, this does not similarly apply to pDCs in mice ^38^. The promoter region of CD317 contains multiple cis-regulatory elements for transcription factors, including STAT ^39,40^. Engagement of STAT induces the expression of CD317 after exposure to type I and type II IFNs ^24–26^, consistent with our *in vitro* data, and lending support to the concept that CD11c^+^CD88^+^ cells in the blood can upregulate CD317 during active EAE under the influence of IFNγ. One ligand of CD317 that has been identified as the surface molecule immunoglobulin-like transcript 7 (ILT7) ^41,42^, which is expressed by immature plasmacytoid DC ^38^. Furthermore, the adhesion of myeloid cells to endothelial cells appears to be partially mediated by CD317 on endothelial cells ^23^, which presumable interact with ILT7 on myeloid cells. Thus, the effects we observed on clinical EAE and the distribution of CD11c^+^CD88^+^CD317^+^ cells after anti-CD317 therapy may be due to binding of anti-CD317 mAB to CD11c^+^CD88^+^CD317^+^cells in blood, or to endothelial cells in the CNS, or both. Our data favor an effect of anti-CD317 on CD11c^+^CD88^+^CD317^+^cells, as the differences in the percentages of CD11c^+^ cells and CD11c^+^CD88^+^ in the blood, brain, and spinal cord in the anti-CD317 treatment group were driven by changes in the percentages of CD11c^+^CD88^+^CD317^+^cells. It was difficult to detect CD11c^+^CD88^+^CD317^+^cells in blood. These observations suggest an expression of CD317 on CD11c^+^CD88^+^ in the blood immediately prior to their egress from the blood into the CNS.

Expression of CD317 by CD11c^+^CD88^+^ cells is partially induced by IFNγ. This was shown through *in vitro* culture of CD11c^+^CD88^+^ cells. This observation provides one possible explanation for the distinct clinical phenotypes observed in active EAE vs adoptive transfer EAE. In active EAE, the activation of CD4^+^ T cells by myelin autoantigen and CFA results in the generation of autoreactive Th17 and Th1 cells, the latter producers of IFNγ. Consequently, a microenvironment is created outside of the CNS that provides exposure to IFNγ that may upregulate CD317 on CD11c^+^CD88^+^ cells in active EAE. However, the adoptive transfer of myelin-reactive CD4^+^ T cells, which includes both Th1 and Th17 cells, provides IFNγ signaling preferentially inside the CNS after the T cells have migrated there ^29–31^. Thus, in the setting of adoptive transfer EAE, CD11c^+^CD88^+^ cells lack IFNγ outside the CNS to induce CD317.

Interestingly, our human CSF transcriptome analyses placed CD11c^+^CD88^+^CD317^+^cells within a population of myeloid cells that was previously defined as CSF microglia ^11^,^32–34^. We found that activated, but not naive murine microglia concomitantly express CD11c, CD88, and CD317. We believe these data indicate that bone marrow-derived CNS CD11c^+^CD88^+^CD317^+^ cells associated with CNS autoimmune disease resemble activated parenchymal microglia. At this point, it is unclear whether these are independent cell populations with similar phenotypes and function, or whether there is a sequential development from one to the other. Future research will focus on determining which of the murine myeloid subsets in EAE recently defined by scRNA-seq express CD317 ^43^.

Pro-inflammatory myeloid cells during the pathogenesis of EAE have been described previously. Along these lines, central roles were reported for CCR2^+^ myeloid cells ^44^, tissue resident classical DC ^45^, microglia vs peripherally seeded monocyte-derived DC ^20^, and pro inflammatory iNOS^+^ macrophages vs anti-inflammatory Arg-1^+^ macrophages ^45^. Our observations add to the complexity of categorizing myeloid cell populations into DC or monocytes/macrophages based on surface markers alone. Our results suggest that detailed morphological and functional profiling is required to provide a more profound categorization and function-based understanding of the role of these cells during different stages of EAE.

In conclusion, here we show that CD11c^+^CD88^+^CD317^+^ cells appear to be critical mediators of disease in EAE that are intimately associated with the onset of disease. These cells originate from peripheral myeloid cells and their disease-propagating effects can be successfully antagonized using a mAb directed against CD317. We speculate that this pathway holds potential as a novel therapeutic option in MS.

## Acknowledgments

Funded by a Merit Review grant (federal award document number (FAIN) I01BX001674) from the United States (U.S.) Department of Veterans Affairs, Biomedical Laboratory Research and Development (O.S.).

---

**Supplementary Figure 1.**
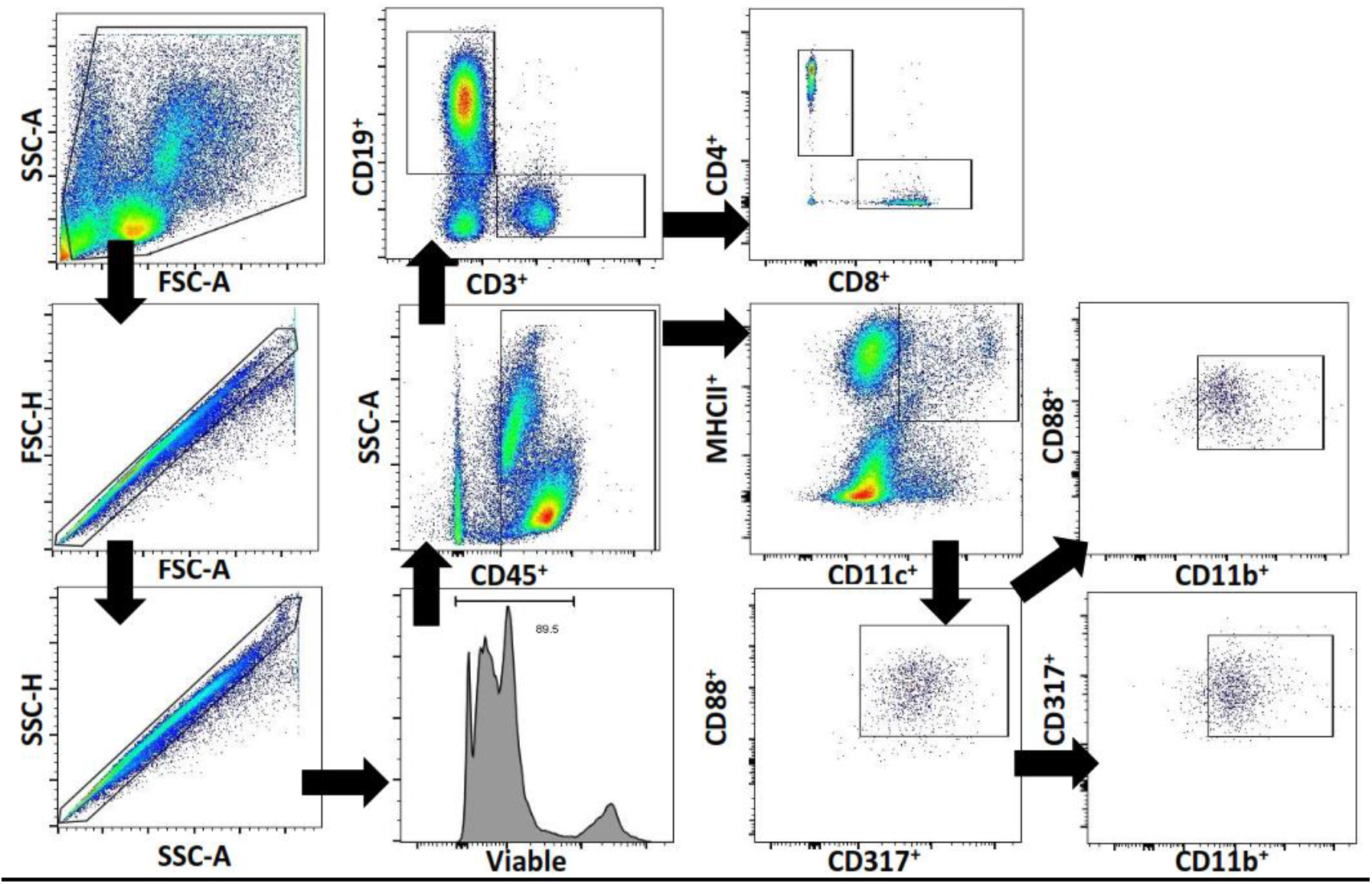
Flow cytometry study gating strategy. Cells were gated according to morphology side scatter (SSC-A) vs forward scatter (FSC-A). Doublets were excluded (FSC-A vs FSC-H and SSC-A vs SSC-H). Live cells were selected using the viability dye. Flow cytometry density plots showing gating strategy used to identify CD11c^+^CD88^+^CD317^+^ cells (CD45^+^MHCII^+^D11c^+^CD88^+^CD317^+^), B cells (CD45^+^CD3^-^CD19^+^), CD4^+^ T cells (CD45^+^CD3^+^CD4^+^) and CD8^+^ T cells (CD45^+^CD3^+^CD8^+^). Each sample contains a minimum of 50 x 10^3^ live events.

**Supplementary Figure 2.**
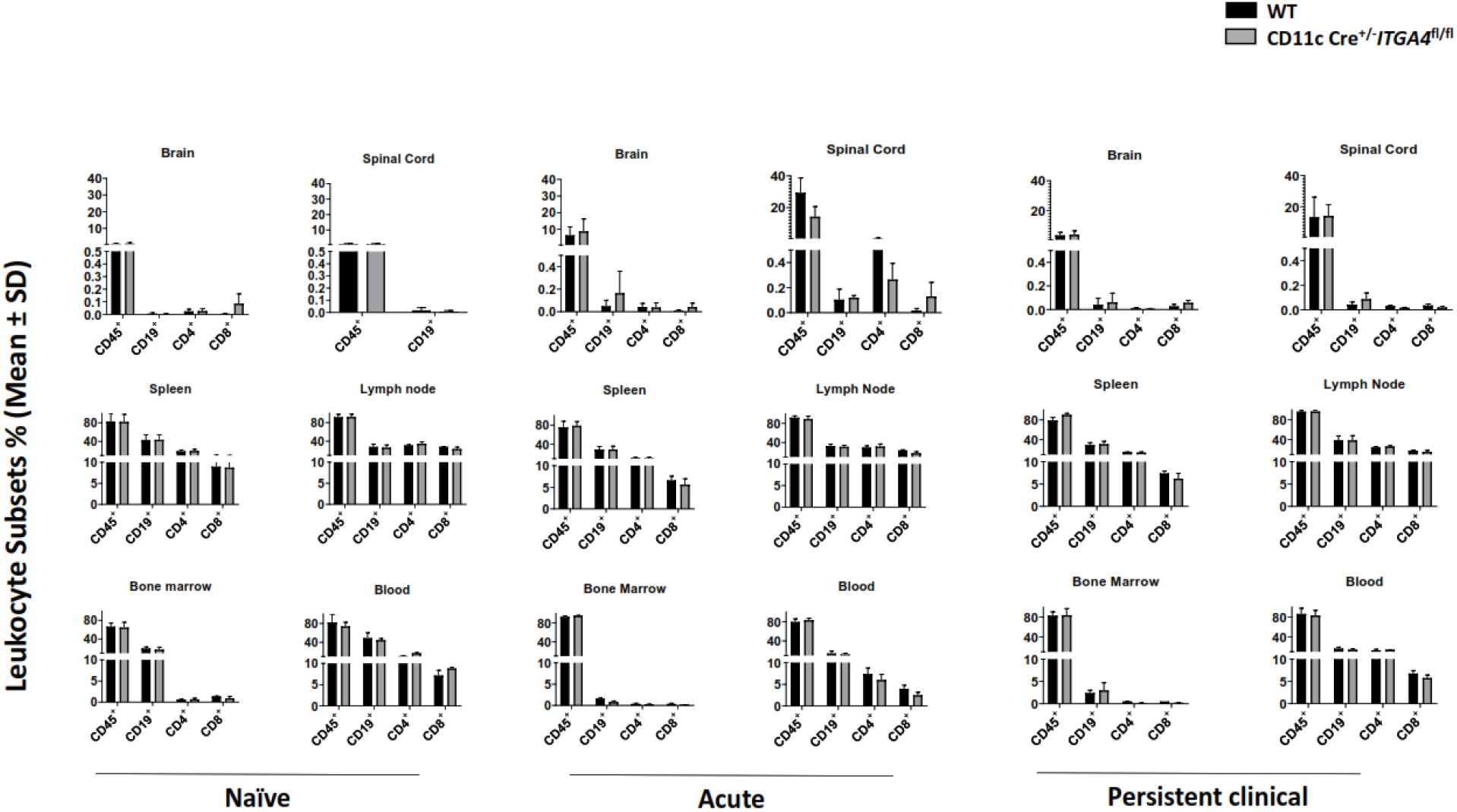
The composition of leukocytes outside the CD11c^+^ lineage is not altered in CD11c.Cre^+/-^*ITGA4*^fl/fl^ mice. The mean ± standard deviation (SD) of leukocyte subsets (%) from total 50 x 10^3^ recorded viable cells in different compartments, including (**A-F**) brain and spinal cord, (**G-L**) spleen and lymph node and (**M-R**) bone marrow and blood, is presented during naïve, acute and persistent clinical actively-induced experimental autoimmune encephalomyelitis (EAE) in CD11c.Cre^+/-^*ITGA4*^fl/fl^ mice or C57BL/6 wild type (WT) controls (N =6 experimental animals per group; data show pooled analysis of all study cohorts). There was no difference in frequency of cell types including CD45^+^, CD19^+^, CD4^+^ and CD8^+^ cells between the two groups during different stages of the disease (P value > 0.05).

